# Cortico-basal white matter alterations occurring in Parkinson’s disease

**DOI:** 10.1101/576991

**Authors:** Bethany. R. Isaacs, Anne. C. Trutti, Esther Pelzer, Marc Tittgemeyer, Yasin Temel, Birte. U. Forstmann, Max. C. Keuken

**Author notes:** Current address: Maastricht University Medical Centre, Department of Experimental Neurosurgery, Room F1.144, P. Debyelaan 25, 6202 AZ Maastricht.

## Abstract

Magnetic resonance imaging studies typically use standard anatomical atlases for identification and analyses of (patho-)physiological effects on specific brain areas; these atlases often fail to incorporate neuroanatomical alterations that may occur with both age and disease. The present study utilizes Parkinson’s disease and age-specific anatomical atlases of the subthalamic nucleus for diffusion tractography, assessing tracts that run between the subthalamic nucleus and *a-priori* defined cortical areas known to be affected by Parkinson’s disease. The results show that the strength of white matter fiber tracts appear to remain structurally unaffected by disease. Contrary to that, Fractional Anisotropy values were shown to decrease in Parkinson’s disease patients for connections between the subthalamic nucleus and the pars opercularis of the inferior frontal gyrus, anterior cingulate cortex, the dorsolateral prefrontal cortex and the pre-supplementary motor, collectively involved in preparatory motor control, decision making and task monitoring. While the biological underpinnings of fractional anisotropy alterations remain elusive, they may nonetheless be used as an index of Parkinson’s disease. Moreover, we find that failing to account for structural changes occurring in the subthalamic nucleus with age and disease reduce the accuracy and influence the results of tractography, highlighting the importance of using appropriate atlases for tractography.

## Introduction

The subthalamic nucleus (STN) is a small region located in the basal ganglia (BG) that is integral to a range of motor behaviors and cognitive functions [1]. Abnormal activity of the STN is implicated in a number of neurodegenerative and neurological disorders including Parkinson’s disease (PD). Here, increased indirect pathway activity is thought to increase the inhibition of motor plans rather than reducing inhibitory control [2]. Accordingly, the STN is a common neurosurgical target for deep brain stimulation (DBS) for PD patients who no longer appropriately respond to pharmacological interventions, where standard targeting is facilitated by the use of MRI and stereotaxic atlases [3].

However, these atlases are often based on a normal population and fail to account for neuroanatomical variability occurring for a variety of reasons, including age and disease [4–9] It is widely acknowledged that the anatomy of the STN varies substantially across healthy individuals, with *in-vivo* size estimates ranging from 50mm^3^ to 270mm^3^ (See [10] and references therein). Additionally, age-related changes associated with the STN show the location in standard MNI space shifts in lateral direction in the elderly population [11–14] with additional alterations of STN volume and location occurring PD [15].

Moreover, the STN demonstrates a complex connectivity profile both within the BG and with the rest of the cortex [16–24]. With regards to PD, both the structural and functional connectivity of the STN has been shown to predict the future outcome and relative success of DBS treatment [25]. This is supported by electrophysiological and functional (f)MRI results which show that specific cortico-basal connections are functionally altered in PD [26–28]. Furthermore, the existing variability in the success of DBS suggests the presence of individual differences in the integrity of specific connections between the STN and different cortical regions.

DBS of the STN is however associated with a number of psychiatric side-effects, cognitive, and emotional disturbances [29,30]. One explanation for these side-effects relates to the somatotopic arrangement of functionally dissimilar cortical projections within the STN [31–34]. In DBS, the implanted electrode may directly stimulate, due to suboptimal placement, or spread current to functionally disparate sub-regions of the nucleus which in-turn interfere with the typical connectivity between the STN and limbic or cognitive cortical areas [35,36].

Given the neuroanatomical alterations that occur in the STN due to orthologic aging or PD, it is crucial to investigate whether additional group specific changes extend to their structural connectivity. The current paper first aims to investigate whether there are disease specific alterations in the connectivity of cortical areas to the subthalamic nucleus in PD patients by using group specific atlases of the STN, and second, to assess whether any connectivity measures may be correlated with disease progression. We chose six cortical areas based on their functional involvements in limbic, cognitive, and motor processes, known to be affected in PD [37–41]. Cortical areas consisted of the pars opercularis of the inferior frontal gyrus (Pop), the anterior cingulate cortex (ACC), the dorsolateral prefrontal cortex (DLPFC), primary motor cortex (M1), supplementary motor area (SMA), and pre-supplementary motor area (pre-SMA). Notably, the we use these results to highlight the importance of using group specific atlases for STN identification when ultra-high field (UHF) MRI is not available, given the scarcity of UHF MRI sites relative to the number of DBS centers [42–47].

## Materials and methods

### Subjects

Seventy PD patients and thirty-one age-matched healthy controls participated in the study (Table 1) (see [48] for more details on subject population). Patients were not required to discontinue their medication for the purposes of this study. The gender imbalance in the PD group was due to the fact that PD is 1.5 times more likely to occur in men than in women [49–51]. Disease related variables were obtained from PD patients, which include UPDRS III scores taken both on and off medication, duration of disease in years, and side of symptom onset (left or right), all obtained from an expert neurologist [52]. Disease progression, as a measure of severity, is calculated by dividing each patients UPDRS off III score by the duration of the disease in years [53]. Medication response is calculated by dividing the UPDRS off III score by the respective UPDRS on [54]. All healthy controls self-reported no history of psychiatric or neurological disease, and PD patients reported no other neurological complaints than PD. The study was approved by the ethical committee of the University Hospital of Cologne, Germany.

**Table 1:** Descriptive statistics.

|  | Parkinson's Disease | Healthy Control |
| --- | --- | --- |
| **Age (years)** | 62.01(8.62) | 61.94(10.21) |
| **Gender** | 54m/16f | 25m/6f |
| **Disease Duration (years)** | 6.51(4.64) | - |
| **UPDRS III on** | 14.46(7.03) | - |
| **UPDRS III off** | 29.84(12.18) | - |
| **Symptom onset (side)** | 33l/37r | - |

The mean (SD) demographic statistics for both the Parkinson’s Disease and healthy control group. UPDRS: Unified Parkinson’s Disease Rating Scale.

### MRI acquisition

Whole-brain anatomical T1-weighted and diffusion-weighted images were acquired for each subject with a Siemens 3T Trio scanner (Erlangen, Germany). T1-weighted images were obtained using a 12-channel array head coil with the following parameters: field of view (MDEFT3D: TR = 1930 ms, TI = 650 ms, TE = 5.8 ms, 128 sagittal slices, voxel size = 1 × 1 × 1.25 mm^3^, flip angle = 18°). dMRI images were obtained via a spin-echo EPI sequence with a 32-channel array head coil (spin echo EPI: TR = 11200 ms, TE = 87ms, 90 axial slices, voxel size = 1.7 mm isotropic, 60 directions isotropically distributed (b-value = 1000 s/mm^2^). Distortions due to eddy currents and head motion were corrected using FSL (Version 5.0; www.fmrib.ox.ac.uk/fsl) [55]. Additionally, to provide an anatomical reference for motion correction, seven images without diffusion weighting (b0 images) were acquired at the beginning and after each block per ten diffusion-weighted images. The diffusion-weighted images were then registered to these b0 images (see [48] for more details regarding the data acquisition).

### Registration

#### MRI

All registration steps were conducted using both linear and nonlinear functions with FLIRT and FNIRT (as implemented in FSL version 5.0). All registrations were performed on skull stripped and brain extracted images. T1 weighted images were first linearly registered to the MNI152 T1 1mm brain template with a correlation ratio and 12 DOF. An additional nonlinear transform was applied using the FNIRT function with standard settings, including the previously obtained affine transformation matrix. Individual T1-weighted scans in native space were registered to the respective no-diffusion (b0) images with a mutual information cost function and 6 DOF. A standardized midline exclusion mask in MNI152 space was registered to each subjects b0 images through multiple transforms, by combining the transformation matrices outputted via previous registrations. The midline exclusion mask was visually checked and realigned with an additional registration if necessary. Each step during the registration process was visually assessed for misalignments by comparing several landmarks (ventricles, pons, corpus callosum, cortical surface).

#### Cortical Atlases

The six cortical areas were obtained from http://www.rbmars.dds.nl/CBPatlases.htm, created with tractography methods, based on both human and non-human primate neuroanatomy [56–58]. The separate cortical masks were extracted from MNI152 1mm space. The cortical atlases were thresholded at 25% to minimize the occurrence of over estimating the region during registration procedures, which were achieved with a nonlinear transform from MNI152 1mm to individual b0 space using the previously generated transformation matrices from the anatomical registrations, with a nearest neighbor interpolation and 12 DOF (see Figure 1).

**Figure 1.**
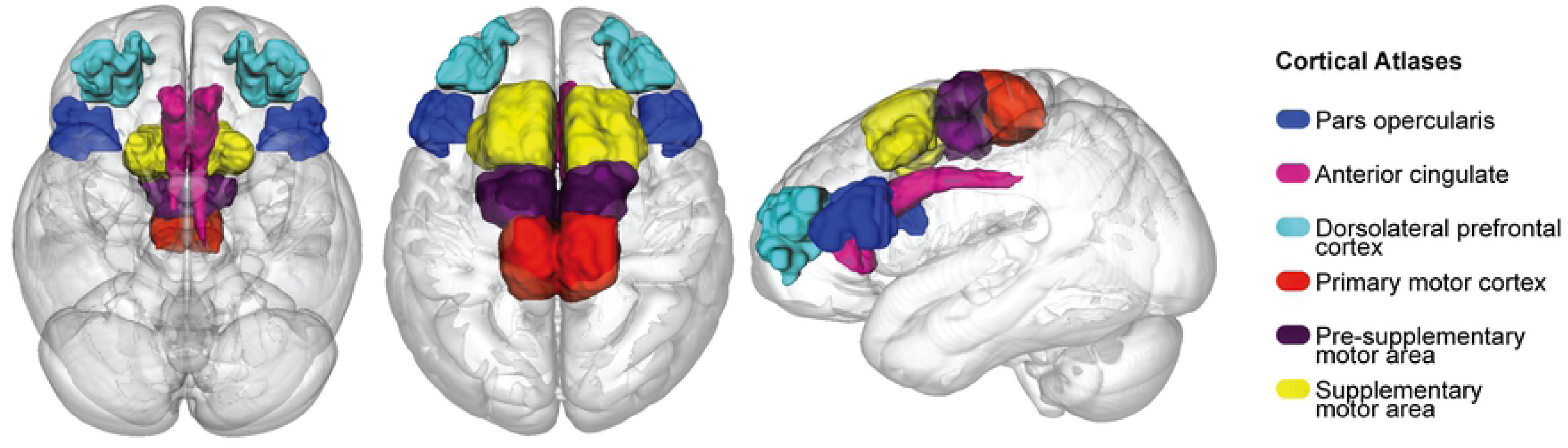
Cortical atlases used for probabilistic tractography and the diffusion tensor models in MNI152 1mm space, which consist of the pars opercularis (POp), anterior cingulate cortex (ACC), dorsolateral prefrontal cortex (DLPFC), primary motor cortex (M1), presupplementary motor area (pre-SMA) and supplementary motor area (SMA).

#### STN Atlases

Group specific PD and elderly probabilistic atlases of the STN were obtained for the respective groups from [59] (Figure 2) (see https://www.nitrc.org/projects/atag_pd/ for probabilistic atlases and ATAG data) and were transformed from MNI152 1mm space to individual b0 space using a nonlinear transform and thresholded by 25%. The non-zero voxel volume in mm^3^ for each atlas was as follows: PD left = 77; PD right = 70.13; HC left = 164.75; HC right = 138.38 and for the center of gravity (CoG) in MNI152 1mm space: PD left x = −10.44, y = −13.04, z = −8.16; PD right x = 11.84, y = −13.18, z = −89; HC left x = −10.56, y = −13.87, z = −7.10; HC right x = 12.10, y = −12.97, z = −6.20.

**Figure 2.**
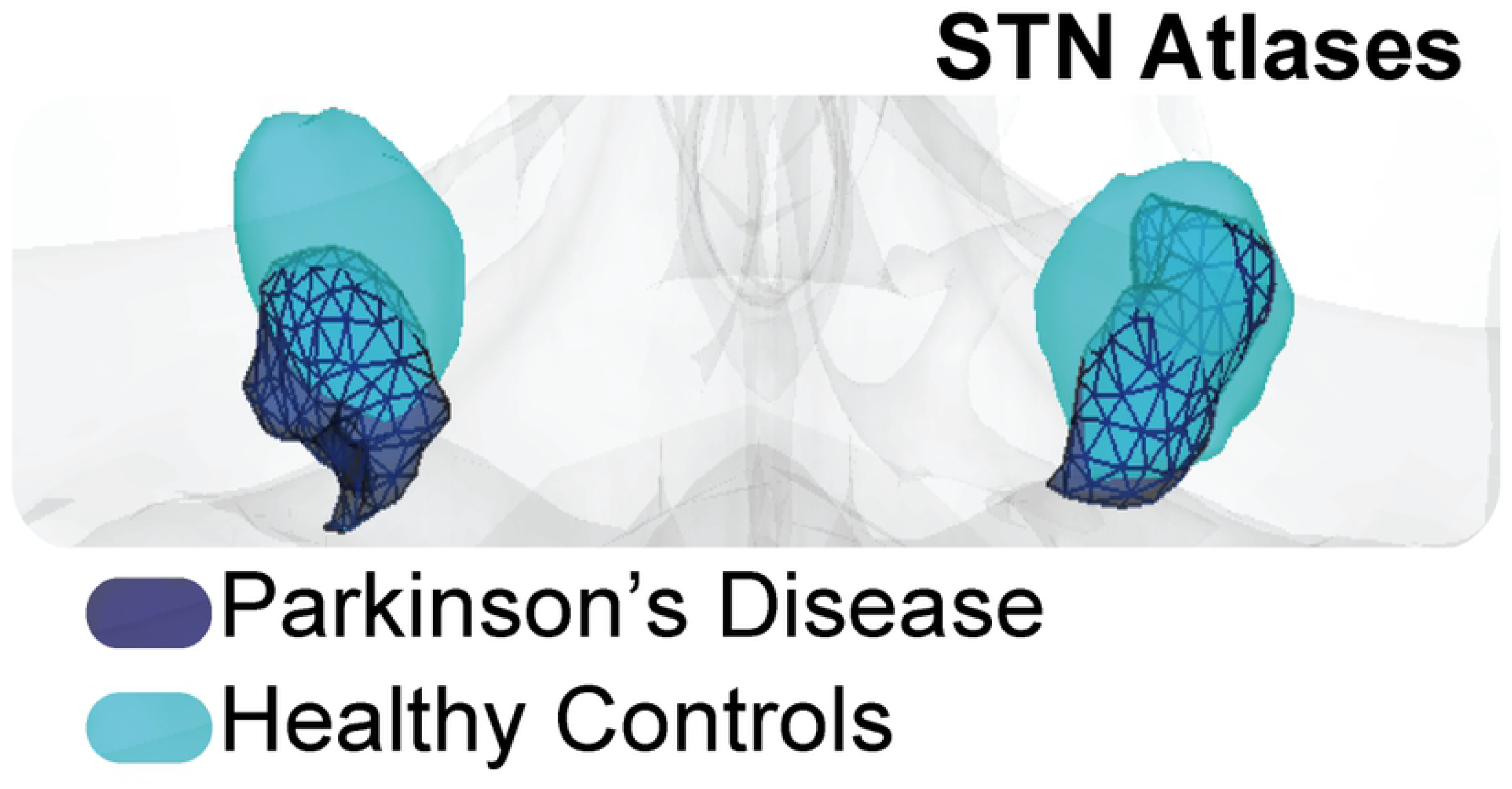
STN atlas in MNI152 1mm space where the PD STN is in dark blue, and the HC in light blue.

### Probabilistic Tractography

Probabilistic tractography was run between *a priori* defined cortical areas and group specific STN’s. Diffusion image preprocessing and analyses were achieved using FSL 5.0. The two most likely diffusion directions per voxel were estimated using the bedpostX function as implemented in the FDT toolbox with standard settings [60]. Subsequently, probabilistic tractography (probtrackX) was conducted to calculate continuous structural connections between the respective seed and target region(s). ProbrackX was run with standard settings (curvature threshold 0.2, 5000 samples, 0.5mm step length, 2000 steps) in each subjects’ native diffusion space, separately for left and right hemispheres and aided by the inclusion of a midline exclusion mask.

The term tract strength here is used to index a probability density function, quantifying the ratio of how many streamlines directly and continuously commence from a seed region and terminate at a target area. This density function is a commonly used measure for inferring the strength of structural white matter tracts [60–62]. For more robust measurements, we created an average of each pair of seed-to-target and target-to-seed streamlines [63,64] To control for spurious tracking, the tracts were thresholded by 10, whereby any voxel containing less than 10 direct samples were excluded from further analyses [64].

We calculated the axial diffusivity (AD), fractional anisotropy (FA), and mean diffusivity (MD) of the seed-to-target and target-to-seed paths derived from the tract strength probability density function approach mentioned in the above section. This was achieved by fitting a voxel wise diffusion tensor model with a weighted least squares regression to each subjects’ diffusion image using the DTIFIT function from FDT. Each FDT path was thresholded so that only paths with at least 75 samples where included for further analysis to yield a conservative anatomical representation. Then each pair of corresponding paths were combined (seed-to-target and target-to-seed), binarized and averaged per hemisphere. From these normalized FDT paths we extracted the AD, FA, and MD values per tract, per subject.

### Statistical Methods

All statistical analyses were conducted within a Bayesian framework (Table 2) using the BayesFactor toolbox [65] in R [66], interpreted in light of the assumptions proposed by [67] and adapted by [68]. To test whether there were any group differences in either tract strength or DTI derived metrics, we used Bayesian ANOVAs. For Bayesian ANOVAs, each BF reported in the output is a ratio of the model’s predictive success relative to a null hypothesis, following a mixed effects JZS Bayesian framework [69,70]. Additionally, where appropriate, we include a comparison between the most likely and the second most likely model, which indicates how much more likely the winning model is given the data compared to the second most likely model. Both subject and hemisphere were added as random factors, accounting for unequal sample sizes. All analysis included default prior scales, and where adjustments of multiplicity are required, prior probabilities of the model are automatically adapted.

**Table 2.**
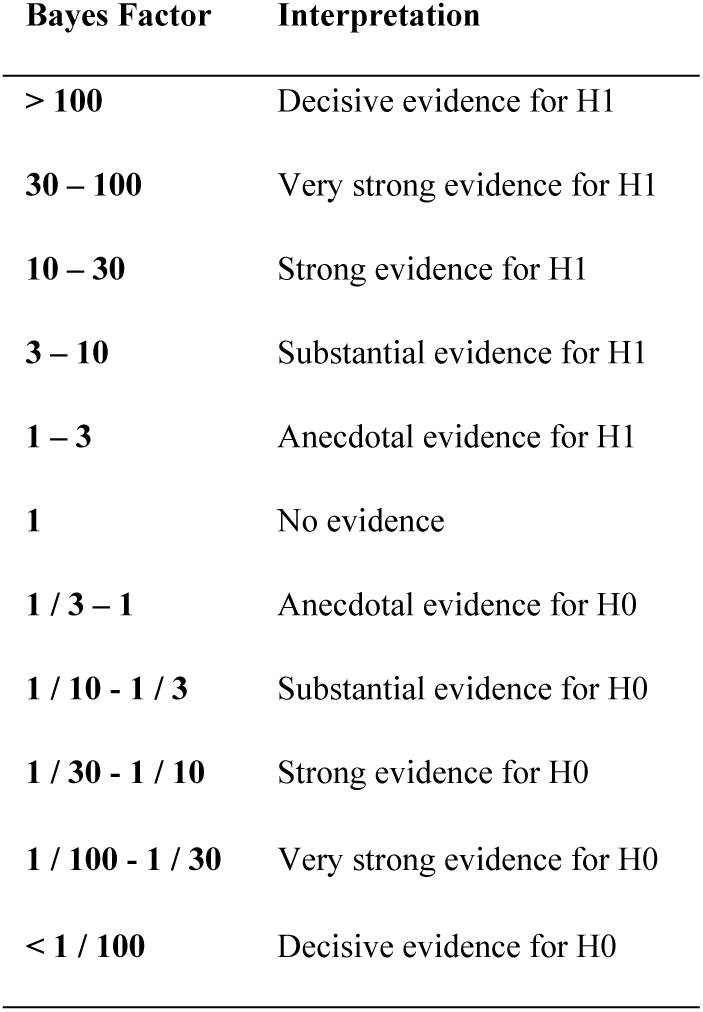
Bayes Factor Interpretation

To test whether disease progression correlated with either tract strength or the DTI derived metrics, we conducted Bayesian correlation analyses in JASP [71]. Disease progression and medication response were used as separate indices of disease severity [72]. All Bayesian tests used a non-informative prior and a medium sized distribution (conjugate distributions on either side).

### Open science

All scripts used to analyse the data can be found at https://osf.io/4uxxs/

## Results

### Group differences between HC and PD

#### Demographics

Two samples Bayesian t-tests were conducted to assess for differences in age and gender across groups. For age, the BF_10_ of 0.23 indicates moderate evidence in favour of the null hypothesis as does a BF_10_ of 0.24 for gender. Therefore, we can assume that there is no difference in gender or age between groups and these variables are not included as covariates for further analyses.

#### Motion Parameters

Additional Bayesian t-tests were conducted to test for differences across groups in each of the directional (x, y, z) translation and rotation parameters, which index how much the subject moves during the MRI. All results were in favour of the null hypothesis (rotation x: BF_10_ = 0.51, rotation y: BF_10_ = 0.33, BF_10_ = 0.25, translation x: BF_10_ = 0.40, translation y: BF_10_ = 0.34, translation z: BF_10_ = 0.49). Accordingly, motion parameters are not included as a covariate in further analyses.

#### Tract Strengths

We first set out to test whether there were differences in tract strength between healthy control subjects and PD patients with a mixed effects ANOVA, incorporating subject and hemisphere as random factors (see Figure 3 and table 3). When using group specific atlases of the STN, the largest model incorporated a main effect of structure and group, without interaction (BF_10_ = 2.61e+175), which is 191 times more likely than the model incorporating an interaction term (BF_10_ = 1.37e+173). This provides decisive evidence for tract strengths varying with both group and structure, but without an interaction.

**Table 3.**
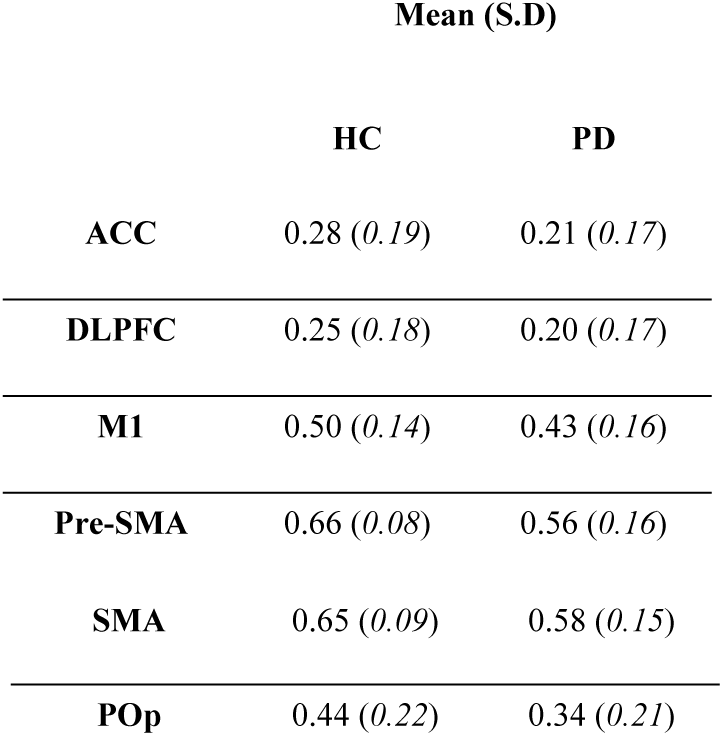
Tract strength descriptive statistics per tract, per group.

**Figure 3.**
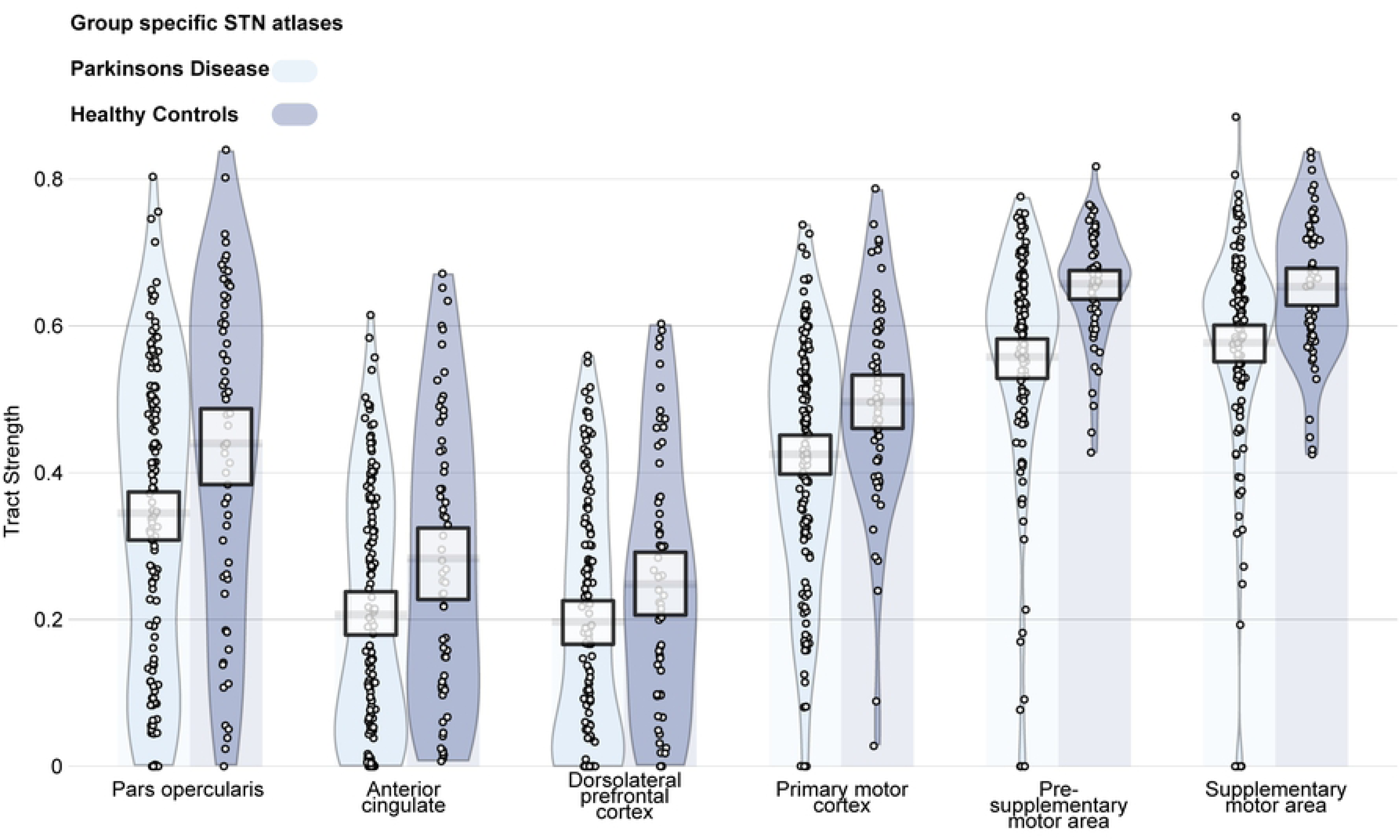
Tract strengths collapsed across hemisphere per structure, with healthy control subjects in purple and PD patients in blue. Tracts are measured from 0 to 1, which is representative of the ratio of the total number of tracts reported between the STN and the given cortical structure. Each point within each element represents a single subject. The width of each element represents the smoothed density. The columns overlapping each bar (each beginning at zero) represent the central tendency, and the bands overlapping each element reflect the 95% highest density intervals.

#### DTI metrics: group differences

To test whether there were group differences in the white matter composition, we extracted the AD, FA, and MD values of the six different tracts. Separate ANOVAs were run to assess AD, FA, and MD across groups (table 4). The model with the most evidence for difference in AD included differences across structure and group, but with no interaction (BF^10^ = 7.92e+148). For MD, the strongest model incorporated only differences across structure (BF_10_ = 5.21e+127).

**Table 4:** Diffusion Tensor Imaging (DTI) descriptive statistics of axial diffusivity, fractional anisotropy and mean diffusivity per tract, per group

|  | Mean (S.D) |  |  |  |  |  |
| --- | --- | --- | --- | --- | --- | --- |
|  | AD |  | FA |  | MD |  |
|  | HC | PD | HC | PD | HC | PD |
| ACC | 1.21e-03<br>(5.21e-05) | 1.18e-03<br>(4.46e-05) | 4.03e-01<br>(2.43e-02) | 3.94e-01<br>(2.85e-02) | 8.30e-04<br>(5.12e-05) | 8.22e-04<br>(5.14e-05) |
| DLPFC | 1.188e-03<br>(4.81e-05) | 1.18e-03<br>(5.47e-05) | 3.67e-01<br>(2.20e-02) | 3.38e-01<br>(2.25e-02) | 8.55e-04<br>(4.45e-05) | 8.74e-04<br>(5.55e-05) |
| M1 | 1.28e-03<br>(7.82e-05) | 1.25e-03<br>(7.54e-05) | 0.41<br>(0.4e-01) | 4.09e-01<br>(3.99e-02) | 8.91e-04<br>(8.62e-05) | 8.91e-04<br>(8.65e-05) |
| Pre-SMA | 1.17e-03<br>(5.01e-05) | 1.16e-03<br>(5.02e-05) | 4.01e-01<br>(2.30e-02) | 3.86e-01<br>(2.58e-02) | 8.23e-04<br>(5.34e-05) | 8.27e-04<br>(5.34e-05) |
| SMA | 1.26e-03<br>(6.74e-05) | 1.27e-03<br>(8.35e-05) | 0.39<br>(0.26e-01) | 3.77e-01<br>(3.23e-02) | 9.16e-04<br>(6.95e-05) | 9.27e-04<br>(9.03e-05) |
| POp | 1.25e-03<br>(6.74e-05) | 1.31e-03<br>(6.06e-05) | 0.35<br>(0.2e-01) | 3.50e-01<br>(2.27e-02) | 9.41e-04<br>(6.26e-05) | 9.37e-04<br>(5.58e-05) |

When assessing FA, decisive evidence was found for the model incorporating an interaction between group and structure with a BF_10_ of 4.57e+184, which is 4.2 times more likely than the second largest model which includes only main effects of structure and group (BF_10_ = 1.06e+184). Post-hoc Bayesian t-tests revealed strong evidence for differences between groups for FA values between STN and POp with a BF_10_ of 18.53, with higher FA values for healthy controls than PD patients. Substantial evidence was found for FA values differing across groups between the STN and the ACC (BF_10_ = 3.05), which are also higher in healthy controls than PD patients. Decisive evidence was found for the DLPFC (BF_10_ = 5.31e+10) and pre-SMA (BF_10_ = 68.31) connectivity profiles, again both with higher FA values for healthy controls than PD patients (Figure 4).

**Figure 4.**
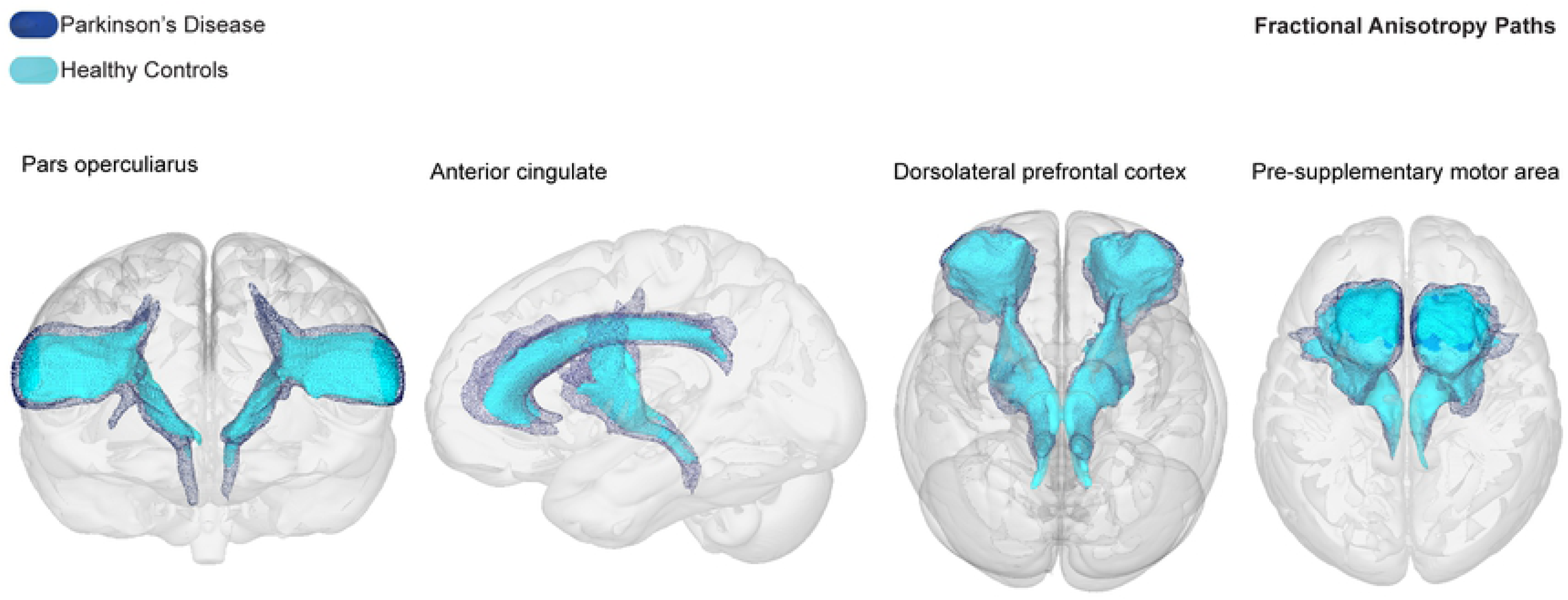
Averaged FA tracts per group running between the STN to the POp, ACC DLPFC, and pre-SMA, with PD tracts in dark blue and healthy control tracts in light blue. In all tracts, the FA was lower for PD compared to healthy controls.

#### Correlations

Bayesian paired correlations with a Pearson’s Rho correlation coefficient was conducted to assess whether for each PD patient, disease progression or medication response correlated with either their tract strength or respective FA measures [71]. Additionally, because the motor related symptoms of PD often begin and continue to exhibit asymmetrically, the side in which symptom onset was first identified (i.e., left or right side of the body) was counterbalanced across hemisphere [73–75]. Symptom onset initiating on the left side of the body was paired with tract strength or FA values arising from the right hemisphere and vice versa for the left hemisphere (contralateral), and a separate correlation test was conducted for those tracts that occur in the hemisphere on the same side as symptom onset (ipsilateral). This was done in order to control for the lateralization effects of both symptom presentation and brain connectivity and to test whether tract strengths can act as an index of symptom severity.

#### Disease progression with tract strength

All results reported substantial evidence for no correlation between tract strengths and disease progression (table 5).

**Table 5:**
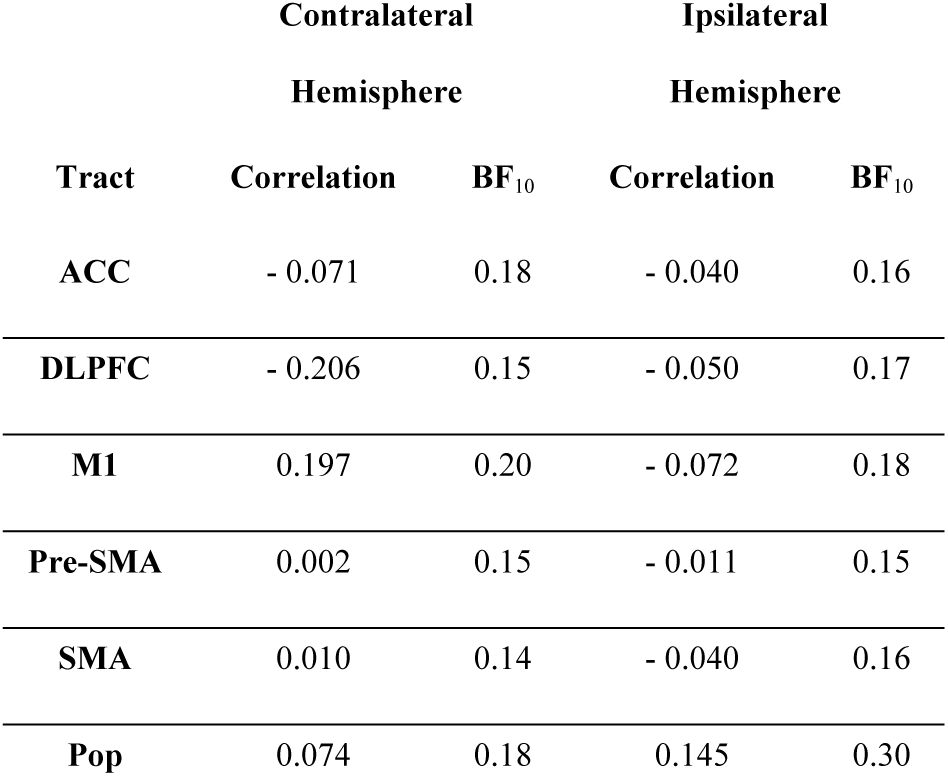
Correlation between disease progression and tract strength

#### Medication response with tract strength

All results reported substantial evidence for no correlation between tract strengths and medication response (table 6).

**Table 6:**
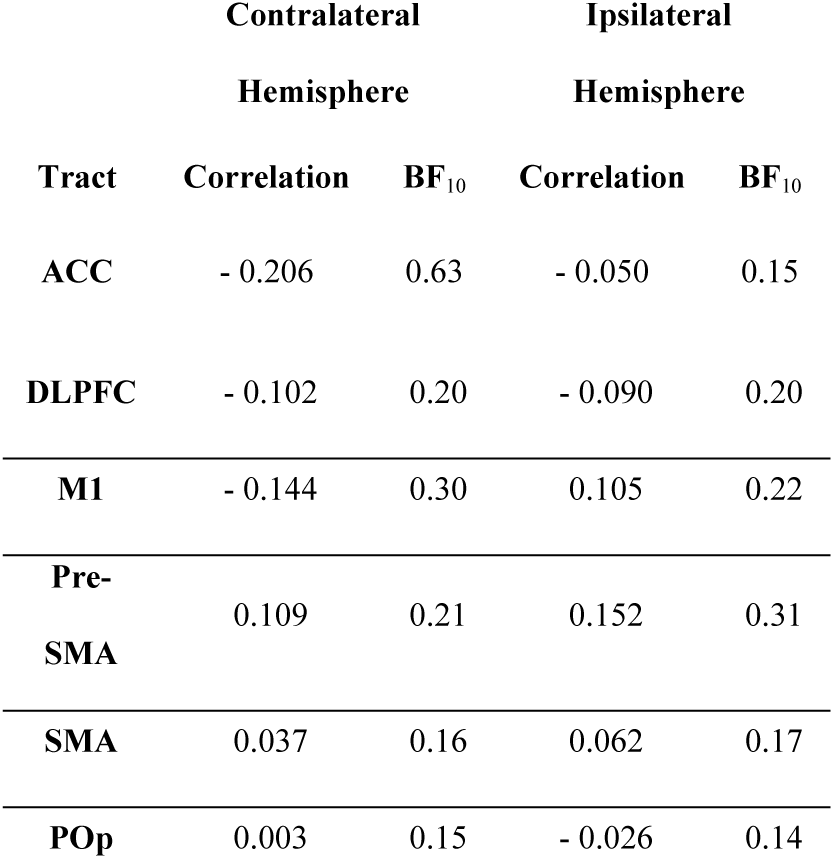
Correlation between medication response and tract strength

| Tract | Contralateral Hemisphere |  | Ipsilateral Hemisphere |  |
| --- | --- | --- | --- | --- |
|  | Correlation | BF<sub>10</sub> | Correlation | BF<sub>10</sub> |
| ACC | - 0.206 | 0.63 | - 0.050 | 0.15 |
| **DLPFC** | - 0.102 | 0.20 | - 0.090 | 0.20 |
| **M1** | - 0.144 | 0.30 | 0.105 | 0.22 |
| **Pre-SMA** | 0.109 | 0.21 | 0.152 | 0.31 |
| **SMA** | 0.037 | 0.16 | 0.062 | 0.17 |
| **POp** | 0.003 | 0.15 | - 0.026 | 0.14 |

#### Disease progression with FA

The only FA path to show strong evidence of a correlation with disease progression was the DLPFC ipsilateral score (r = 0.364, BF_10_ = 16.50), where side of symptom onset and hemisphere were the same. All other results reported either anecdotal or substantial evidence for no correlation between FA and disease progression (table 7).

**Table 7:** Correlation between disease progression and fractional anisotropy

| Tract | Contralateral Hemisphere |  | Ipsilateral Hemisphere |  |
| --- | --- | --- | --- | --- |
|  | Correlation | BF | Correlation | BF |
| **ACC** | 0.141 | 0.28 | 0.214 | 0.71 |
| **DLPFC** | 0.204 | 0.61 | 0.364 | 16.50 |
| **M1** | - 0.027 | 0.14 | - 0.009 | 0.15 |
| **Pre-SMA** | 0.111 | 0.23 | 0.09 | 0.18 |
| **SMA** | - 0.026 | 0.14 | -0.003 | 0.15 |
| **POp** | 0.161 | 0.36 | 0.106 | 0.22 |

#### Medication response with FA

All results reported substantial evidence for no correlation between FA and medication response (table 8).

**Table 8:** Correlation between medication response and fractional anisotropy

| Tract | Contralateral Hemisphere |  | Ipsilateral Hemisphere |  |
| --- | --- | --- | --- | --- |
|  | Correlation | BF | Correlation | BF |
| ACC | 0.125 | 0.25 | 0.030 | 0.14 |
| DLPFC | 0.100 | 0.21 | 0.018 | 0.14 |
| M1 | 0.037 | 0.16 | 0.084 | 0.19 |
| Pre-SMA | 0.017 | 0.24 | 0.105 | 0.22 |
| SMA | 0.014 | 0.15 | 0.041 | 0.16 |
| POp | 0.055 | 0.17 | - 0.042 | 0.16 |

## Discussion

The current study assessed the strength and microstructural changes occurring in predefined connectivity profiles between the STN and motor, limbic, and cognitive related cortical areas between PD patients and healthy elderly age-matched controls using group specific atlases of the STN.

### PD Disease specific alterations

For all six cortical areas, the tract strength was lower for the PD group. Moreover, none of the tract strengths between the STN and the cortical areas correlated with measures of disease progression or medication response. It therefore appears unlikely that the strength of any of the measured tracts may be used as a biomarker for PD. However, the STN for both groups showed strongest structural connections to the motor cortices, which likely reflects the role of the STN in motor control.

Diffusion tensor models were applied to draw quantitative measurements of each white matter tract. For the original analysis, we found evidence for a reduction in FA for the STN to POp, ACC, DLPFC, and pre-SMA tracts in PD patients compared to healthy controls. The POp is situated anterior to the premotor cortex and has been implicated in motor inhibition [76] which is referred to as the ability to suspend a premeditated motor response to a stimulus or an ongoing response [77]. It has also been proposed that the POp is the origin of “stop signal” behaviors, whereby the inhibition of a motor response results from direct stimulation of the subthalamic nucleus [78]. Moreover, the primary STN-ACC circuit functions to monitor behaviors that involve conflict and therefore task switching and changing decisions [79–81]. Given the symptomatic profile of PD patients, it seems plausible that the STN-POp and ACC connectivity profiles would be structurally and or functionally effected by the disease [82,83].

Relatedly, associated functions of STN-pre-SMA circuit also include response inhibition [84], action choices [85–87], task switching and internally generated movements [88,89], which are shown to be disrupted in PD. Assuming structure both shapes and constrains function [90–92], compromised white matter tracts indexed by increased diffusivity and reduced FA could result in abnormal functioning and lead to clinically overt behaviors [93]. A dysfunctional STN-pre-SMA circuit could result in parkinsonian symptoms including micrographia, dysarthia, bradykinesia and hypokinesia, all of which involve a lack appropriate action selection, timing, and irregular task switching [94–96]. A dysfunctional STN-DLPFC circuit could reflect impaired motor control, PD related cognitive decline, and affective complaints [97–99] as well as being linked to dopaminergic abnormalities [100]. However, while reduced FA in specific STN-cortical circuits could be utilized as a biomarker for PD, it is difficult to infer the exact biological mechanisms underlying alterations in diffusion metrics relative to disease. FA has been considered as a summary measure of white matter integrity, that is highly sensitive to microstructural changes, but less sensitive to the type of change [101–104], though theoretically, a reduction in FA could be driven by a singular or combination of altered AD, MD, or radial diffusivity.

Moreover, white matter consists not only of axons, but oligodendrocytes, astrocytes and microglia and therefore structural changes can affect any of these properties, each of which is associated with a different function [105,106]. Studies have shown that FA correlates with myelination which is associated with speed conduction, though this is dependent on the formation and remodeling of oligodendrocytes and differentiation of oligodendrocyte precursor cells (OCPs) whose function is to determine the production, length and thickness of internodes and therefore also likely to contribute to the FA signal [107–110]. Fewer studies have assessed diffusion parameters in relation to astrocytes, though their contribution to FA signals is likely to be significant given their large occupying volume within both grey and white matter [109,111].

Physiologically, a disruption or structural abnormality occurring anywhere along the axon, for example due to changes in myelination, impaired astrocyte propagation or suboptimal OCP proliferation and differentiation, would impede the rate of conduction and transmission between structures and consequently result in functional impairments [112]. Additionally, more widespread changes in myelin and internode plasticity can be driven by region-specific mechanisms [113,114]. In the case of PD, local signals arising from dopaminergic cell loss with the substania nigra, or the pathological hyperactivity of the STN could drive the observed structural changes in cortico-basal white matter connections. However, due to the complex timeline and microscopic spatial resolution of these neurochemical and anatomical changes, it is currently not possible to identify which process corresponds with in-vivo human dMRI based FA measures.

Further, diffusivity has been correlated with partial voluming effects arising from free-water [115]. Free-water reflects the presence of water molecules that are not restrained by cellular barriers and therefore do not show a preference for direction, which may be increased in the presence of cellular damage [116]. Thus, the presence of free-water may influence biases on diffusion metrics which can result in a reduction in FA and or an increase in MD [117,118]. For instance, free-water present in diffusion has been shown to reflect FA changes occurring in other PD affected areas such as the substantia nigra [119,120]. Additionally, the measure of tract strength was taken via a probability density function (PDF), which despite being shown as a robust assessment, remains controversial. Measurements indexing for instance, relative strength via dynamic causal models offer a viable alternative [121].

### Correlates of PD disease severity

Overall, we found no evidence for any correlation between either tract strengths or FA values with disease progression or medication response. With one exception, we found a positive correlation for FA values within the STN-DLPFC connectivity profile increasing with disease progression when the side of symptom onset was matched with hemisphere. An increased FA indicating restricted diffusion along a single direction is not necessarily compatible with explanations of neurodegenerative processes when assuming a higher FA implies increased myelination and axonal density which usually decrease with disease progression. It may be possible that the increased FA is explained by an attempted compensatory, neuroplasticity mechanism and or functional reorganization rather than a direct neurodegenerative process [122,123], or a response to atypical dopaminergic modulation and levodopa intake [124–126]. Such an adaptive reorganization of structural and functional pathways would, however, occur long before the onset of clinical symptoms, which is not in line with the rather progressed stage of the PD population within this study [73,75]. We therefore remain speculative as to the explanation of this result.

### Considerations

The use of MRI poses several challenges when imaging small subcortical nuclei such as the STN [127]. In the current study, the resolution of the anatomical and diffusion sequences was rather large when considering the size of the STN [128]. Imaging the STN is subject to partial voluming effects and blurring of the voxels near the borders of the nucleus, which may contain different tissue types and or fiber bundles of neighboring structures ^24,129^. This is further complicated by probabilistic atlases being inherently larger than is often anatomically exact and require registration between template and native space. Such registration procedures employ simple scaling factors that can fail to optimally incorporate morphometric and densitometric variability between individuals [130] which can in turn affect the accuracy of subsequent analysis. We account for this by using group specific atlases, thresholding atlases, and incorporating both rigid and affine transformations during registration procedures. In the supplementary section we included a number of additional analysis to investigate the effects of atlas accuracy.

Manual segmentation of the both the STN and cortical areas for all individuals would be the golden standard, however, the data in the current study did not allow for manual parcellation of the STN or of structurally distinct cortical areas [131,132]. Relatedly, the visualization of the STN would benefit from the use of sub-millimeter resolution imaging with UHF MRI and/or susceptibility-based contrasts [13,133].

Lastly, we do not assess for gender differences. While sexual dimorphisms in PD have been reported [50,96,134,135], it remains controversial as to how sensitive standardized scores such as the UPDRS are at identifying gender differences [134,136]. In addition, we include a relatively small sample size with an unbalanced male to female ratio.

## Conclusions and future directions

To conclude, the strength of white matter tracts within the hyper-direct pathway appear unaffected by the pathophysiology of PD. However, decreased FA values of the STN-POp, STN-DLPFC and STN-pre-SMA tracts may be used as a biomarker for disease, though the exact biological mechanisms driving these disease specific alterations in FA remain elusive. Regardless, the differences we find are in the connections to cortical areas involved in preparatory motor control, task monitoring and decision making, rather than cortical areas governing motor output. Further, the results indicate that it is recommended to use an atlas that accounts for anatomical changes associated with PD rather than only age matched controls. See the supporting information for a control analysis to support the use of group specific atlases. Future work should focus on the use of higher field strengths, alternative tractography methods and harmonization of techniques used to investigate PD [137,138]. Until then, we show that using atlases that are specific to your population can aid analysis where UHF MRI and or manual segmentations are not possible.

Tractography methods hold great promise for their contribution to identification of disease, differential diagnoses between subtypes of parkinsonian syndromes and the application of DBS [139,140]. Such applications require assessment of the biological foundations of diffusion metrics and neuroanatomical factors with specific subsets of disease scales used to evaluate presence and severity.

## Supporting information

**S1 File: Supporting information**: supplementary materials

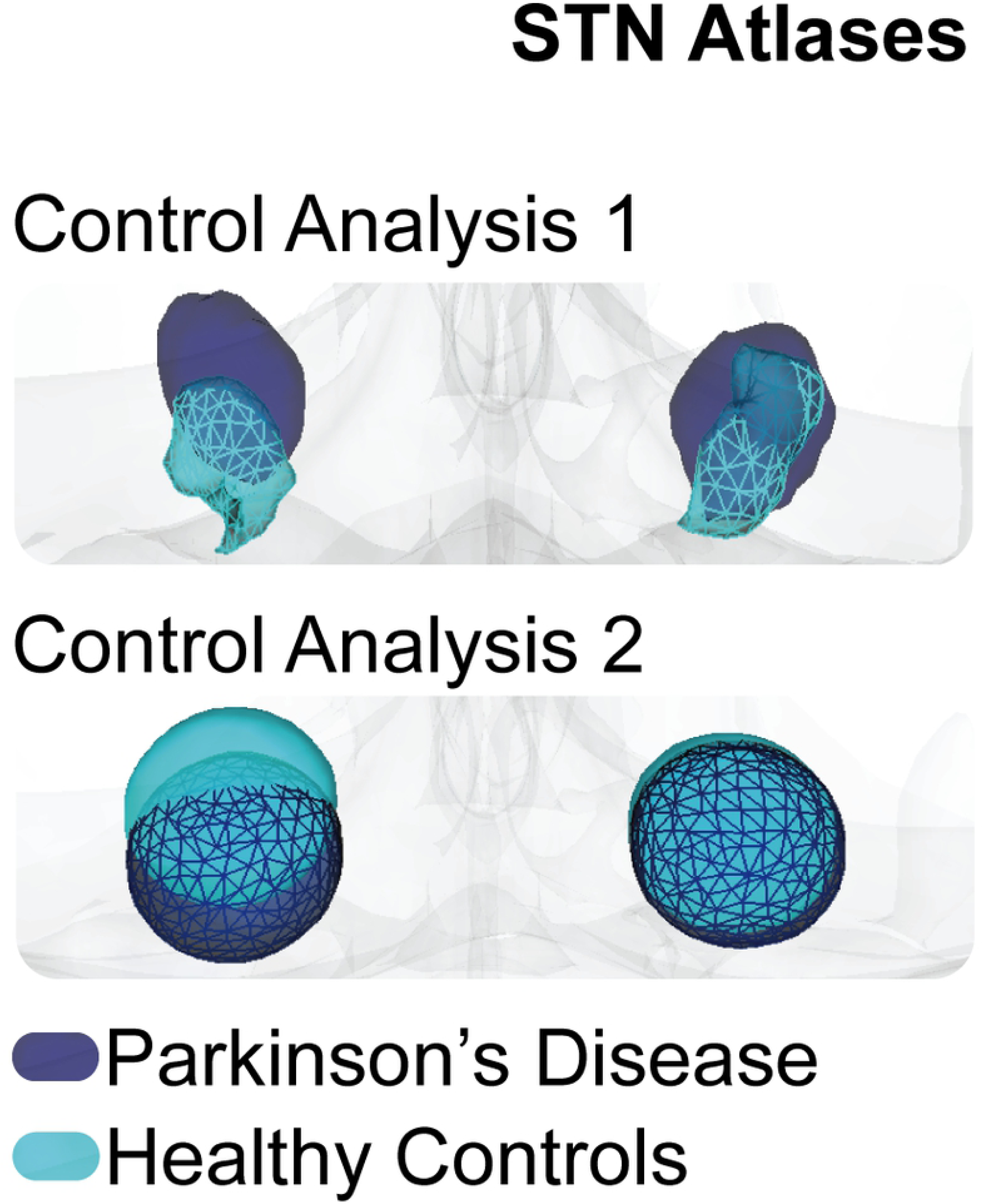

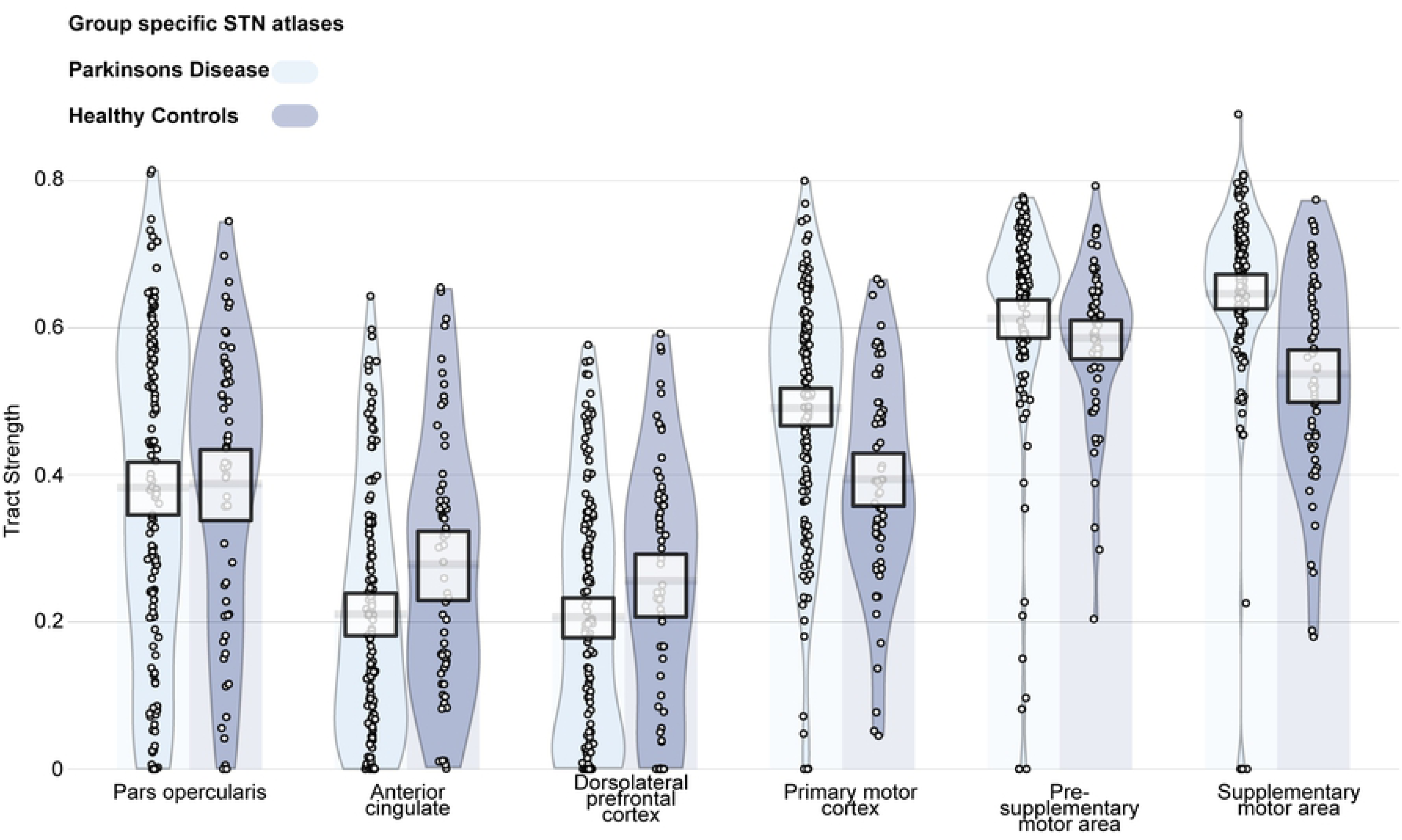

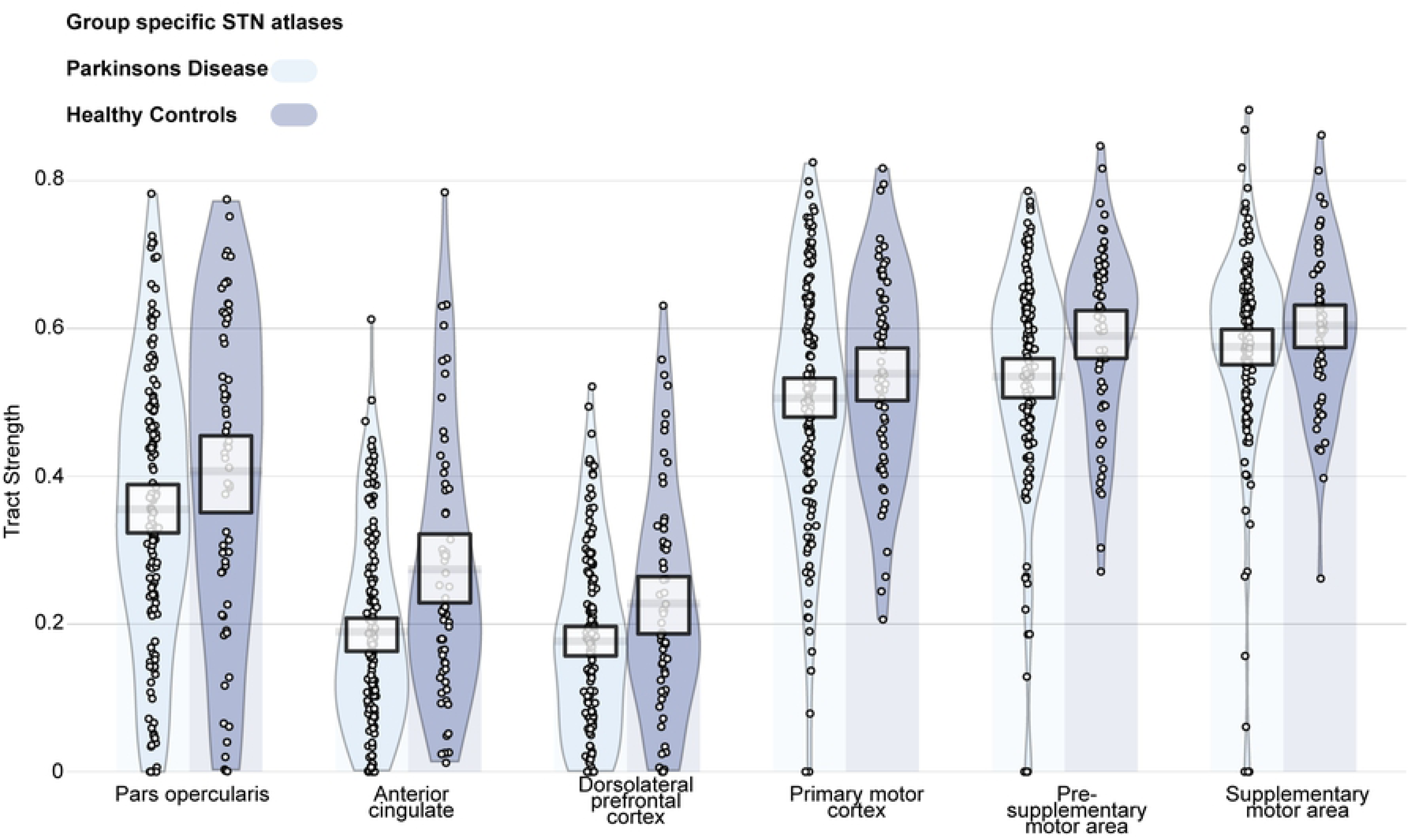

